# Frequency and duration of sensory flicker controls astrocyte and neuron specific transcriptional profiles in 5xFAD mice

**DOI:** 10.1101/2024.05.20.594705

**Authors:** Sara Bitarafan, Alyssa F. Pybus, Felix G. Rivera Moctezuma, Mohammad Adibi, Tina C. Franklin, Annabelle C. Singer, Levi B. Wood

**Affiliations:** Parker H. Petit Institute for Bioengineering, Georgia Institute of Technology; George W. Woodruff School of Mechanical Engineering, Georgia Institute of Technology; Wallace H. Coulter Department of Biomedical Engineering, Emory University and Georgia Institute of Technology; School of Biological Sciences, Georgia Institute of Technology

**Keywords:** Flicker, audiovisual sensory stimulation, Alzheimer’s disease, transcriptional profiling, WGCNA, glia, neurons

## Abstract

**Background:** Current clinical trials are investigating gamma frequency sensory stimulation as a potential therapeutic strategy for Alzheimer’s disease, yet we lack a comprehensive picture of the effects of this stimulation on multiple aspects of brain function. While most prior research has focused on gamma frequency sensory stimulation, we previously showed that exposing mice to visual flickering stimulation increased MAPK and NFκB signaling in the visual cortex in a manner dependent on duration and frequency of sensory stimulation exposure. Because these pathways control multiple neuronal and glial functions and are differentially activated based on the duration and frequency of flicker stimulation, we aimed to define the transcriptional effects of different frequencies and durations of flicker stimulation on multiple brain functions.

**Methods:** We exposed 5xFAD mice to different frequencies of audio/visual flicker stimulation (constant light, 10Hz, 20Hz, 40Hz) for durations of 0.5hr, 1hr, or 4hr, then used bulk RNAseq to profile transcriptional changes within the visual cortex and hippocampus tissues. Using weighted gene co-expression network analysis, we identified modules of co-expressed genes controlled by frequency and/or duration of stimulation.

**Results:** Within the visual cortex, we found that all stimulation frequencies caused fast activation of a module of immune genes within 1hr and slower suppression of synaptic genes after 4hrs of stimulation. Interestingly, all frequencies of stimulation led to slow suppression of astrocyte specific gene sets, while activation of neuronal gene sets was frequency and duration specific. In contrast, in the hippocampus, immune and synaptic modules were suppressed based on the frequency of stimulation. Specifically,10Hz activated a module of genes associated with mitochondrial function, metabolism, and synaptic translation while 10Hz rapidly suppressed a module of genes linked to neurotransmitter activity.

**Conclusion:** Collectively, our data indicate that the frequency and duration of flicker stimulation controls immune, neuronal, and metabolic genes in multiple regions of the brain affected by Alzheimer’s disease. Flicker stimulation may thus represent a potential therapeutic strategy that can be tuned based on the brain region and the specific cellular process to be modulated.

## Introduction

Pharmaceutical strategies targeting a single pathological hallmark of Alzheimer’s disease (AD), such as amyloid beta (Aβ) accumulation have remained challenging, suggesting that successful treatment methods must target multiple key aspects of the disease (1-5). Multiple studies, including ours, have shown that 40Hz visual and/or audio stimulation induced microglial changes associated with engulfment phenotypes with the ability to clear Aβ (6-9). Moreover, we have recently shown that exposing wild-type mice to different frequencies of visual stimulation (20Hz, 40Hz, random) elicited distinct patterns of activation of the nuclear factor kappa-light-chain-enhancer of activated B cells (NFκB) and mitogen activated protein kinase (MAPK) pathways (6), which are central regulators of gene expression, cell survival, and immune function in Alzheimer’s disease (10, 11). Thus, non-invasive audiovisual stimulation holds the potential to simultaneously affect multiple aspects of brain function with different effects depending on the frequency and duration of stimulation, which may provide essential utility for treating multifactorial diseases such as Alzheimer’s disease. Yet, we lack a comprehensive understanding of the effect of different frequencies and durations of combined audiovisual flicker on brain function in the context of Alzheimer’s disease pathology. Given the apparent multi-potency of flicker stimulation, the goal of the current study was to utilize transcriptomics to holistically define the effects of differing frequencies and durations of flicker stimulation on a multitude of brain functions in a mouse model relevant Alzheimer’s disease.

Therapeutic avenues for AD need to address multiple aspects of tissue pathology, going beyond the traditional pathological hallmarks such as Aβ accumulation and neurofibrillary tangles, to assess the effects of therapy on dysregulation of tissue metabolism, microglial-mediated immune dysfunction, impaired astrocyte support function, and reduced neurogenesis among others (12, 13). Collectively, the prior body of work around audiovisual flicker stimulation shows that it has the capability to change important aspects of brain function in multiple settings (6-9, 14). Signaling pathways modulated by flicker stimulation, including the NFκB and MAPK pathways, control numerous aspects of brain function, such as neuronal survival, synaptogenesis, microglial activation and functional state, astrocyte reactivity, among many others (11, 15, 16). We previously showed that specific durations of flicker stimulation induce phosphorylation of these pathways, while different frequencies of flicker have different effects on immune-modulatory cytokine expression (6). Although NFκB and MAPK pathways are master regulators of downstream gene expression that control diverse cellular functions in brain, there remains a gap in knowledge on how different frequency and duration of flicker modulate gene expression that controls neuronal and glia functions.

Here we sought to holistically define the potential multi-potency of flicker stimulation with different frequencies and durations of stimulation in the context of Alzheimer’s disease relevant pathology. To do so, we exposed the 5xFAD amyloidosis mouse model to a range of flicker frequencies (constant light, 10Hz, 20Hz, 40Hz) for durations of 30min, 1hr, or 4hr, then used bulk RNA sequencing (RNAseq) to profile transcriptional changes in the visual cortex (VC) and hippocampus (HIP). We used combined audiovisual stimulation for this study because multimodal stimulation is hypothesized to affect more brain regions and prior studies report this stimulation modulates neural activity and AD pathology in the hippocampus (14, 17). We used weighted gene co-expression network analysis (WGCNA) (18) to identify modules of highly correlated co-expressed genes (**Methods**). We identified 12 modules of genes in the VC and 15 modules of genes in the HIP, many of which were enriched in biological processes related to neuronal or immune function. In the VC, we found all frequency of stimulation elicited fast activation of immune-related genes and slow suppression of synaptic genes. In contrast, within HIP, the flicker driven response was differentially regulated based on frequency of stimulation. Specifically, 4hr of both 10Hz and 20Hz suppressed neural differentiation and development and neuro-signaling genes. While 10Hz caused fast suppression of neuronal differentiation and development genes, 20Hz triggered upregulation of mitochondrial, metabolism and synaptic translation genes. Interestingly, in both regions, we found that flicker stimulation differentially activated modules that are enriched in a specific cell type in a frequency and duration dependent manner. In visual cortex, an astrocyte enriched module was differentially regulated by different frequencies and durations while neuronal enriched modules were regulated by all frequencies in a duration dependent manner. In contrast, within HIP, neuronal enriched genes were mainly mediated by frequency of stimulation not duration. Our finding that different frequencies and durations of stimulation distinctly affect neuronal and glial gene modules emphasizes the potential for flicker to target multiple brain cells and brain functions important to Alzheimer’s disease. Our data therefore supports the idea that targeted flicker therapies may be designed to stimulate these functions and provide a new multi-potent avenue of AD therapeutics.

## Materials and Methods

All resources with available Research Identifiers (RRIDs) are listed in Table S11.

### Animal Studies and Sample Collection

All animal work was approved by The Georgia Tech Institutional Animal Care and Use Committee. Male 3–7-month-old 5xFAD on a BL6/SJL background and wild-type littermates were used for this study. Mice were individually habituated for 1hr in a cage before stimulation.

### RNA Sequencing

RNA transcriptomics were quantified from male 3–7-month-old 5xFAD and WT mice exposed to either 10 Hz, 20 Hz or 40Hz audiovisual flickering frequency for a duration of 0.5h, 1h and 4h (**Table S1**). RNA was isolated from visual cortices and hippocampi tissues using Qiagen RNeasy kit (217804; Qiagen; Hilden, Germany) according to the manufacturer’s protocols. Molecular Evolution core at Georgia Institute of Technology sequenced paired ended mRNA using NovaSeq 6000 Sequencing System to obtain a sequencing depth of 30-40 million reads per sample. Prior to sequencing, quality control was performed using a bioanalyzer to determine that the RNA Integrity Number (RIN) of the samples (i.e., >7). A NEBNext Poly(A) mRNA Magnetic Isolation Module and NEBNext Ultra II Directional RNA Library Prep Kit were used to prepare sequencing libraries. RNA alignment was performed by Molecular Research Lab (https://mrdnalab.com) and data was validated for fastq integrity and quality. Four technical replicates of each sample were merged followed by using DNAstar ArrayStar. Next, Qseq reads were mapped using the mouse reference genome GRCm39 (GCF_000001635.27) from the reference database. For the read assignment, the threshold was set at 20bp and 80% of the bases matching within each read. Duplicated reads were eliminated, genes with less than 10 raw counts at least in 25% of the samples were removed from analysis. The non-coding genes were removed from the analysis. All gene counts were normalized using *DESeq2* package in R. To adjust the data for the effects of experiment day, linear mixed model using *removebatcheffect()* function from *limma* packaged was used.

### WGCNA

Weighted gene co-expression network analysis (WGCNA) was conducted in R using *WCGNA* package. WGCNA threshold power of 6 was chosen as it was the smallest threshold the resulted in a scale-free R^2^ value greater than 0.8. Single was constructed in a single block using *blockwiseModules()* using the following parameters: power of 6, a minimum module size of 50, a maximum module size of 12000, a merge cute height of 0.15, correlation type as bicor, and a “mean” TOM denominator. Significant module eigengenes (ME) scores between groups were assessed using linear mixed model using *limma* package available in R.

### Differential abundance analysis

Differential abundance analysis was conducted using *DESeq* package from R programming language. Both frequency and batch were used as variables in the design matrix. Significant level and fold change were computed using *DESeq()* function. Fold change =>0.25 and p<0.05 were considered as significant DEGs and displayed as volcano plot using *EnhancedVolcano()* function in R.

### Gene Ontology

Gene ontology was conducted for each of MEs identified by WGCNA in both regions using PANTHER overrepresentation test on PANTHER 18.0 on the Gene Ontology resource (https://geneontology.org/). Mouse reference genome mm10 GO biological processes complete annotation set was used with Fisher’s exact test to compute significant of the gene sets for each ME.

### Cell type enrichment analysis

Cell type enrichment (CTE) was conducted using protocol from Johnson et. al for each module identified in WGCNA. Briefly, cell type markers from 5 different brain cells (i.e., neuron, microglia, astrocyte, oligodendrocyte and endothelial) were identified from mouse brain proteome and transcriptome studies and were used as reference. Unsupervised enrichment algorithm, *gsva* () function from *gsva* package in R, was used to calculate enrichment score for each cell type across all MEs.

### Custom astrocyte Gene Sets Variation Analysis

To study functional impaction of changes within specific cell types, gene set variation analysis (GSVA) was conducted referencing previously published neuron or astrocyte custom gene sets (19). GSVA is an unsupervised enrichment algorithm which identifies variations of pathway activity by defining enrichment score for gene sets which each contain a set of genes that share same cellular function. Using the R programming language and *gsva* package available on Bioconductor, we identified enrichment of neuron or astrocyte gene sets within neuronal or astrocytic module eigengenes.

### Data analysis and Visualization

Data was analyzed and figures were generated in RStudio (Boston, MA, USA) using the R programming language. Figures were then further polished using Adobe Illustrator (San Jose, CA, USA) or BioRender (Toronto, ON, Canada). Software packages and functions described in this section are denoted by italic font. Heatmaps were generated using the R package *heatmap3*, line graphs and regression plots were created using the packages *ggplot2*. Clustering was conducted using the *hclust* function of the stats package in R using Euclidean distance with Ward. D2 agglomeration method. Outlier detection was conducted in R by calculating Mahalanobis distance of each point from the data’s centroid within each region (Hippocampus, Visual Cortex). Two Hippocampi and one visual cortex samples were detected as an outlier and those samples were omitted prior to analysis. For statistical testing in modules and gene sets, a linear mixed model from *Limma* package available on R programming language was used to determine the group differences (Light vs 10 or 20 or 40 Hz) between each Module defined by WGCNA or gene sets by GSVA. For all analysis, p < .05 was considered statistically significant. All module scores were z-scored.

## Results

### 40Hz vs 20Hz flicker stimulates immune genes in 5xFAD mice

Having previously found that 1hr of 40Hz of visual stimulation activated immune signaling proteins compared to 20Hz stimulation in wild type mice (6), we began the current study by asking if we could identify related transcriptional signatures in 5xFAD mice exposed to 1hr of 40Hz or 20Hz of audiovisual flicker. We exposed male 5xFAD mice and wild-type (WT) littermates to 20 or 40Hz audiovisual flicker for 1hr, then conducted bulk RNAseq of the visual cortex (VC) and hippocampus (HIP) (**Fig. 1A, Methods, Table S1**). Of the 11,312 genes remaining after filtering (**Methods**), we identified 175 differentially expressed genes (DEGs) in the VC (86 up-regulated in 20Hz and 89 up-regulated in 40Hz) and 242 DEGs (68 up-regulated in 20Hz, 174 up-regulated in 40Hz) in the HIP (**Fig. 1B, 1C**). To gain insight into the functional implications of upregulated DEGs, we performed an overrepresentation Gene Ontology (GO) analysis, which revealed significant enrichment of immune-related GO terms in both regions (**Fig. 1D, 1E**). While only 20 DEGs (including *Slc7a1, Cyp1b1, and Alx4,* **Table S2**) are shared across both regions, many of the biological processes enriched in 40Hz in both regions were linked to immune processes. The same analysis on tissues from WT littermates (6) also identified DEGs in both brain regions, but did not identify any significant GO terms, suggesting fewer synchronized transcriptional changes across regions in WT mice at the whole-tissue level (**Fig. S1A**). Nevertheless, our findings in 5xFAD mice indicate that 40Hz audiovisual flicker stimulation may enhance brain immune signaling in the context of amyloid beta pathology. These transcriptional changes are consistent with prior studies showing that 40Hz flicker alters microglial function and signaling and further shows that immune genes in the hippocampus are activated after just one hour of stimulation.

**Figure 1:**
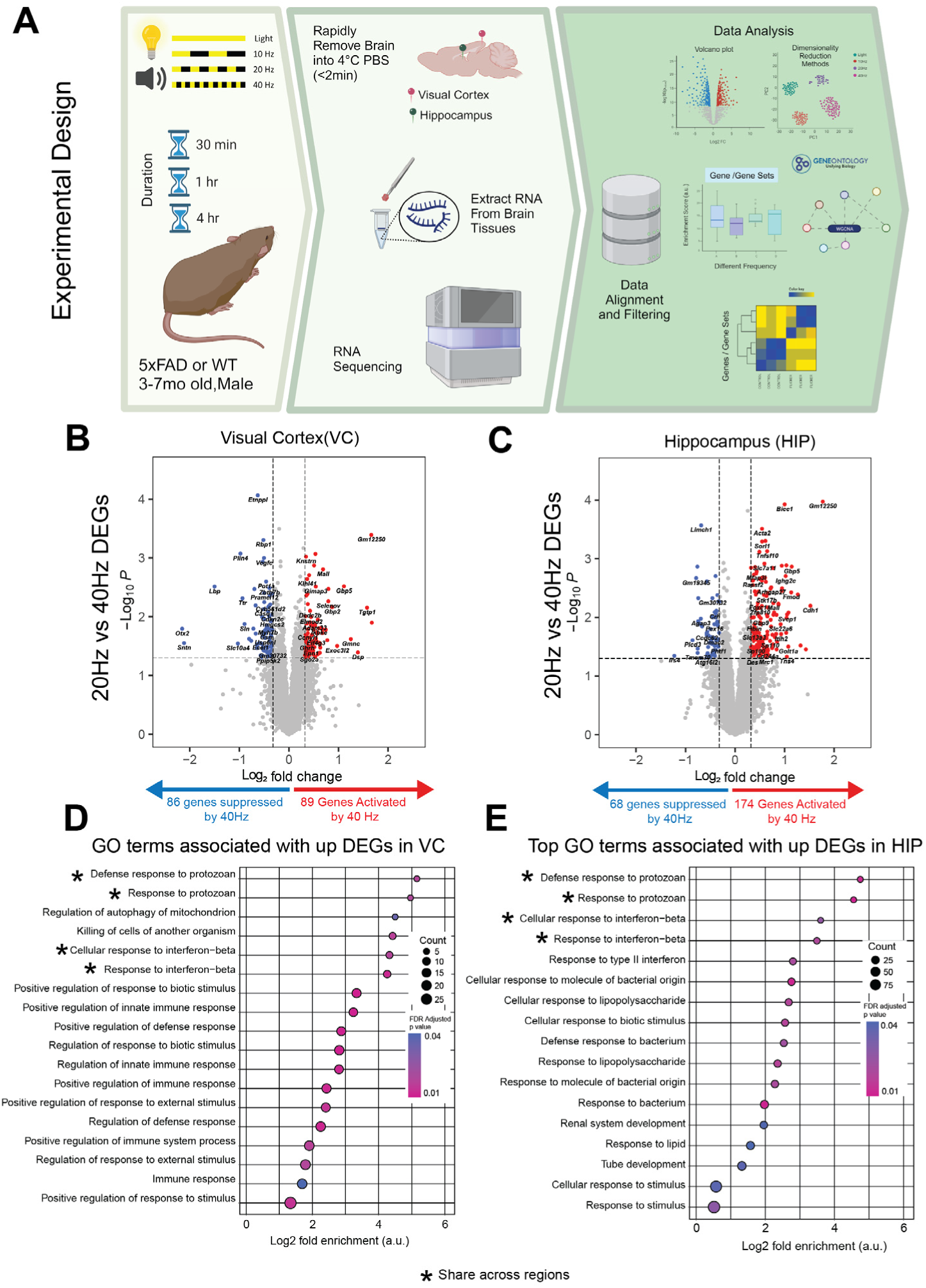
1hr of 40Hz vs 20Hz stimulation stimulates genes associated with immune response in Visual Cortex and Hippocampus of 5xFAD mice. **(A)** Experimental design for audio/visual stimulation. After stimulation, the visual cortices and hippocampi were isolated and sequenced via bulk RNA sequencing followed by statistical analysis (See Supplementary table 1 for statistical details). **(B)** 1 hr of 40Hz or 20Hz stimulation resulted in differentially expressed genes in the visual cortex and **(C)** hippocampus with upregulated genes in red and down regulated genes in blue. Genes of interest are listed in red and were upregulated (log_2_ fold change > 0.25, unadjusted p<0.05). **(D)** Gene ontology analysis of VC genes up-regulated in 40Hz identified significant GO terms associated with immune processes (FDR adjusted p<0.05, dot size represents gene counts and dot colors indicate p-values). **(E)** GO analysis of HIP genes up regulated in 40Hz identified significant terms associated with immune processes (FDR adjusted p<0.05, dot size represents gene counts and dot colors indicate p-values).

### All flicker frequencies stimulated rapid activation of neuroimmune genes and slow suppression of synaptic genes in the visual cortex

We next asked if there were patterns of genes that similarly respond to all frequencies of stimulation in the visual cortex. To holistically understand the data, we employed weighted gene co-expression network analysis (WGCNA) to identify modules of co-expressed genes that were highly correlated across frequency and duration of stimulation (**Methods**). WGCNA identified 12 module eigengenes (MEs) in the VC (**Fig. 2A**), enabling us to collapse the 11,312 genes into 12 modules of highly correlated genes. For genes within each WGCNA module, GO functional annotation (**Methods**) revealed a diverse major functional classification among MEs (**Fig. 2B**). The normalized z -score of each module eigengene (ME) was computed to compare across samples (**Figs. 2C, S2**). The heatmap of all z-scored module eigengenes suggested differences in the ME score patterns in response to different frequencies or durations of stimulation. Next, referencing previously published cell-type specific mouse brain transcriptomes (20, 21), we computed enrichment scores of genes within each MEs via GSVA with high specificity in neurons, microglia, astrocytes, oligodendrocytes, and endothelia (**Methods**). Differing cell-type enrichment indicates differences in cell-type specification for each ME for example VC-ME9 is enriched in astrocyte specific genes while VC-ME6 demonstrated enrichment in neuronal specific genes (**Fig. 2D**). Lastly, to ensure the age of the mice was not a driver of the transcriptional signatures we found here, we correlated age all identified modules (**Fig. S6**) and excluded any that were significantly correlated.

**Figure 2:**
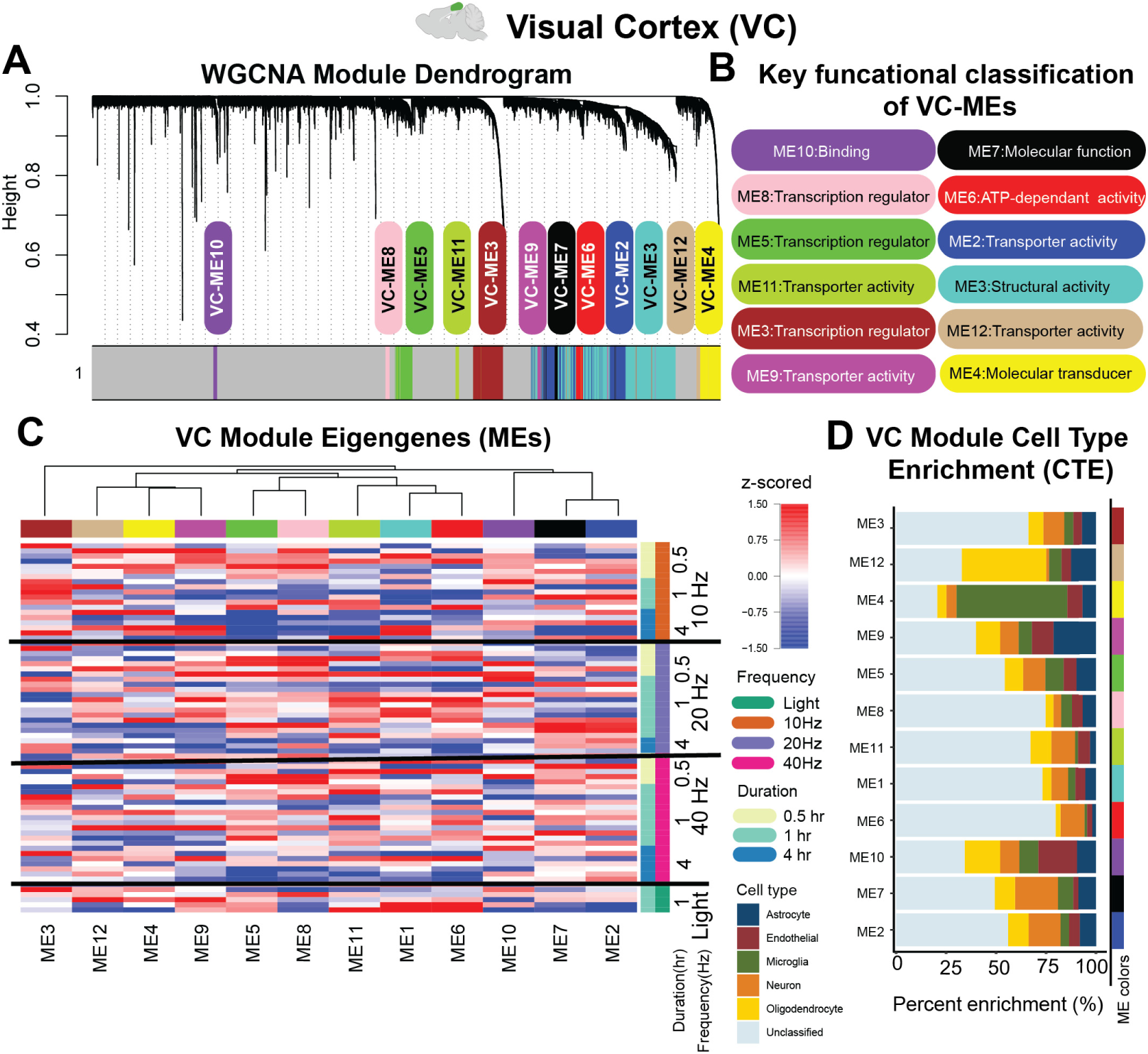
Visual cortex WGCNA identifies modules of co-expressed genes associated with frequency and duration of stimulation. (**A**) WGCNA identified 12 Module Eigengenes (MEs) in VC with each color indicating a different ME. (**B**) Key functional classification of modules were identified via GO Panther function classification. (**C**) Z-scored values for 12 MEs in VC (clustered via Euclidean distance). (**D**) Cell type percent enrichment of all 12 MEs in VC were calculated by crossing-referencing module genes using GSVA against list of genes previously determined as enriched in neurons, oligodendrocytes, astrocytes, microglia and endothelial (Methods). Some MEs were highly enriched by genes expressed by astrocytes (ME9), and neurons (ME6, ME7, and ME2) over other cell-types.

Among all visual cortex MEs, VC-ME8 and VC-ME5 showed consistent changes with duration of flicker, regardless of frequency (**Figs. 3, S2**). VC-ME8 genes were rapidly up-regulated after exposure to any frequency of flicker, then subsided back to Light level by 4hr (**Fig. 3A**). This module did not show notable enrichment for any of cell types assessed in this study (**Fig. 3B**). Interestingly, hub genes of VC-ME8 (**Fig. 3C**) such as *Csnk1a1, Ubn2, Elapor2* and *Dlg4* are all involved in modulating disease progression in many neurodegenerative disease (22-26), including AD. GO analysis of genes comprising VC-ME8 revealed terms associated with regulation of interleukin 17 (IL-17) and type I– interferon (IFN)-mediated signaling (**Fig. 3D**), both of which are associated with brain immune function(27-29). VC-ME8 was also significantly enriched for GO terms associated with hormone signaling and RNA splicing, suggesting a broader effect on brain signaling and transcriptional machinery. In contrast to the rapid activation of VC-ME8, VC-ME5 showed a pattern of slow suppression over duration (**Fig. 3E**) and was not enriched in any specific cell type (**Fig. 3F**). GO processes enriched in VC-ME5 genes are mainly related to synaptic activity and dendrite development (**Fig. 3H**). VC-ME5 hub genes, including *Tmem29*, *Snap29* and *Irs2,* are involved in controlling neuronal excitability, vesicle-mediated transport, and brain immune signaling, which are all dysregulated in AD (30-32) **(Fig. 3G)**. Together, modules VC-ME8 and VC-ME5 indicate that all frequencies stimulate short-term immune function gene expression and slowly suppress gene expression associated with synaptic activity.

**Figure 3:**
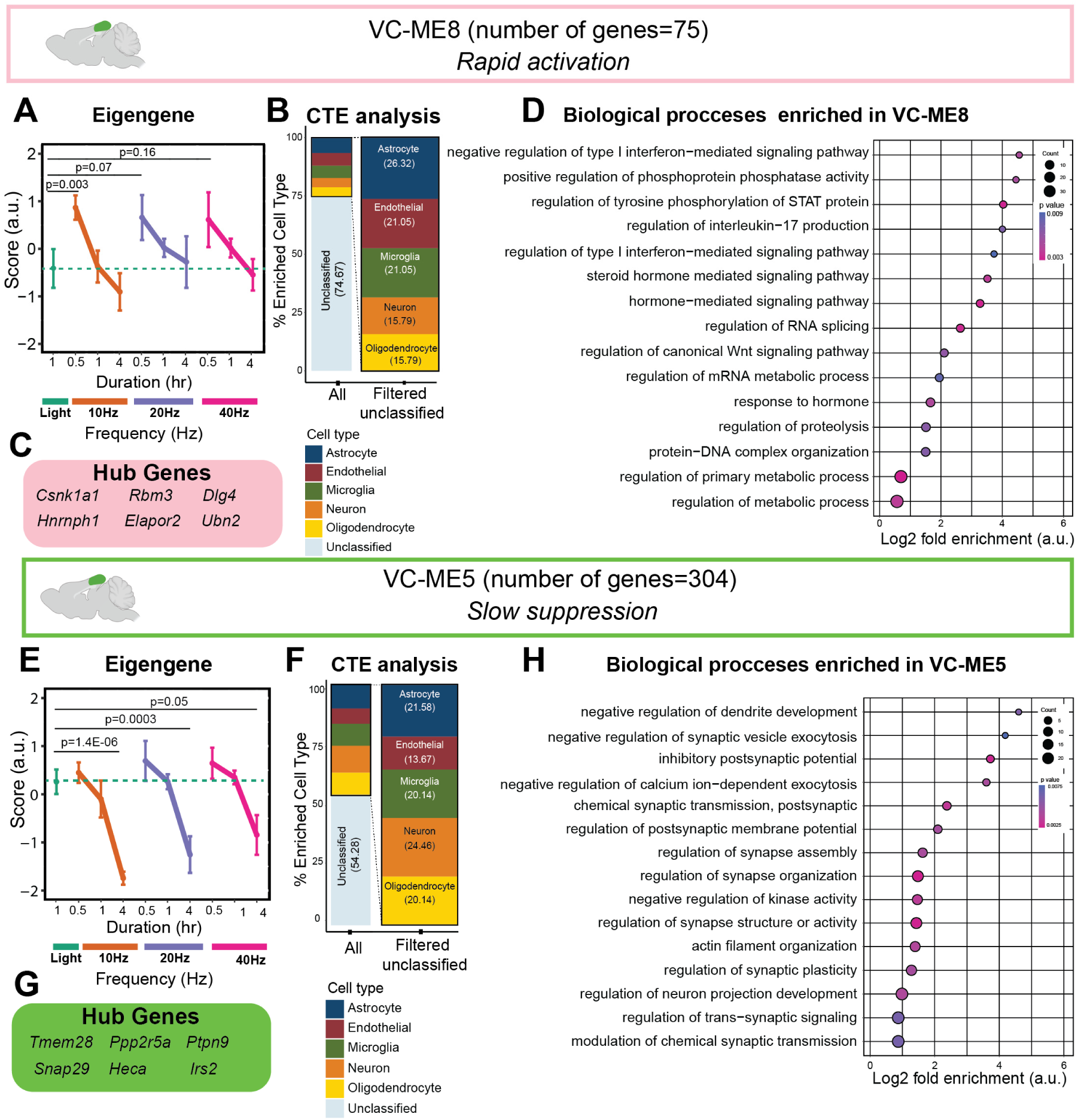
All frequencies of flicker drive rapid activation of neuroimmune genes and slower modulation of synaptic genes in visual cortex. (**A**) VC-ME8 (Pink) genes demonstrated rapid activation in response to all frequencies of stimulation compared to light group (**Table S1**, mean±SEM, linear mixed model). (**B**) Cell type enrichment analysis (CTE) (**Methods**) revealed similar % enrichment in VC-ME8 genes across all cell types (**C**) The six most central genes in the module were determined as hub genes of VC-ME8. (**D**) Gene ontology enrichment of biological processes for VC-ME8 genes (Fisher’s exact test p <0.05) identified neuroimmune changes associated with this module (Dot size represents gene counts and dot colors indicate p-values). (**E**) All frequencies of flicker resulted in slow suppression of VC-ME5 (Green) genes compared to the light group (**Table S1**, mean±SEM, linear mixed model). (**F**) Like VC-ME8, VC-ME5 genes were not highly enriched in any of the 5 cell types assessed here. (**G**) The six most central genes in the module were determined as hub genes of VC-ME5. (**H**) Gene ontology enrichment of biological processes for VC-ME5 genes (Fisher’s exact test p <0.05) identified synaptic and neuro-development changes associated with this module (Dot size represents gene counts and dot colors indicate p-values).

### Astrocyte-enriched genes modulate neuro-signaling is a frequency-duration dependent manner in the visual cortex

Having found shared effects of flicker stimulation among all frequencies, we next looked for visual cortex MEs with distinct patterns of activation in response to different frequencies of flicker stimulation. Stimulation at all flicker frequencies suppressed VC-ME9 genes at either 1hr or 4hr of stimulation (**Fig. 4A**). Interestingly, cell type enrichment analysis revealed that the nearly 40% of VC-ME9 genes are astrocytic (**Fig. 4B**). Astrocyte enrichment is consistent with the appearance of the astrocytic hub gene *S100a1* (**Fig. 4C**), and significantly enriched GO terms related to synaptic regulation which is primarily mediated by astrocytes in the brain (**Fig. 4D**). Because VC-ME9 was astrocyte enriched, we next assessed how different astrocyte-mediated functions are affected by flicker stimulation within VC-ME9 genes by using gene set variation analysis (GSVA) together with our previously published astrocyte-specific gene sets (19)(**Methods**). Specifically, we computed enrichment scores of nine cortical astrocyte gene sets including astrocytic lipid metabolism, astrocytic carbohydrate metabolism, astrocytic protein metabolism and presynaptic astrocyte processes (PAPs) (**Figs. 4E, S4, Table S9**). Interestingly, we found slow suppression of PAPs and lipid metabolism gene sets and rapid activation of carbohydrate metabolism gene sets in response to all frequencies of stimulation (**Fig. 4E**). Collectively, astrocyte specific gene expression changes by flicker stimulation are regulated by both duration and frequency of stimulation in VC of 5xFAD mouse model.

**Figure 4:**
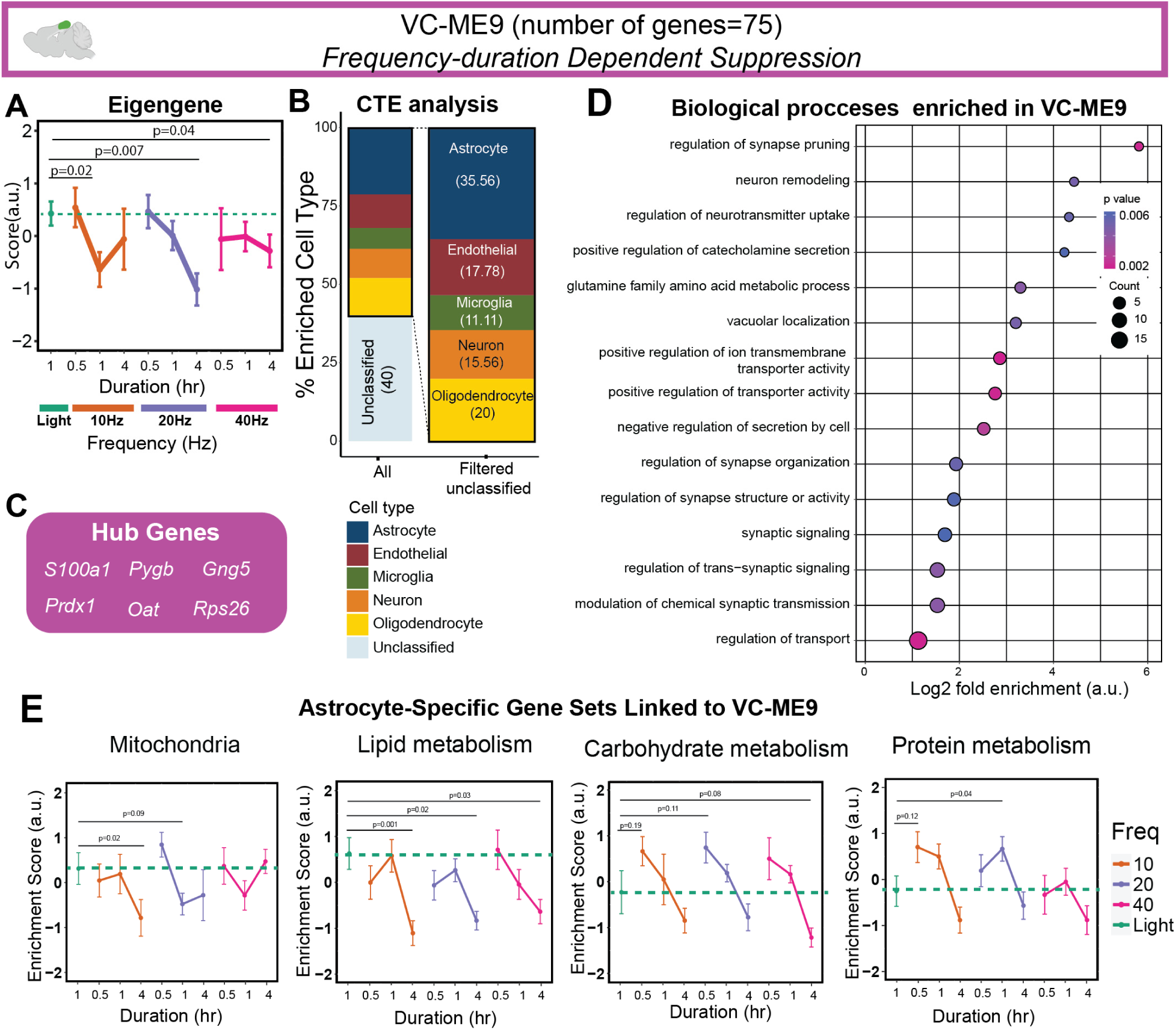
Flicker modulates astrocytic genes that regulate neuro-signaling with frequency and duration-specific effects of simulation in visual cortex. (**A**) VC-ME9 (Magenta) genes demonstrated frequency- and duration-specific suppression in response to flicker (**Table S1**, mean±SEM, linear mixed model). (**B**) Astrocytic genes were the most prominent genes enriched in VC-ME9 (**C**) The six most central genes in the module were determined as hub genes of VC-ME9. (**D**) Gene ontology enrichment of biological processes for VC-ME9 (Fisher’s exact test p <0.05) identified functions associated with synaptic signaling and neurotransmitter activity in this module (Dot size represents gene counts and dot colors indicate p-values). (**E**) Gene set variation analysis of VC-ME9 genes using custom cortical astrocyte specific gene sets (**Methods**) revealed frequency and duration dependent changes in astrocyte-specific functions in response to audiovisual flicker.

### Different frequencies modulate neural functions in a duration-dependent manner in the visual cortex

Since sensory flicker is believed to act via neurons, we concluded our visual cortex analysis by asking if we could identify a neuronally enriched gene signature that responded to flicker stimulation. VC-ME6 was highly enriched (59.4% of classified genes) for neurons and was suppressed across all frequencies (**Fig. 5A-B**). Interestingly, hub genes of VC-ME6 are mainly transcriptional regulators, such as *Dnaja2* and *Vsp35* (33, 34), as well as nervous system development regulators, such as *Pfn2* (35). Of interest, *Dnaja2* is also shown to play protective role against tau aggregation in AD (34, 36, 37)and *Vsp35* gene has shown to be associated with late onset Parkinson disease (38). GO terms associated with VC-ME6 genes were mainly involved in gene expression and transcription regulatory processes (**Fig. 5D**). Like astrocytes, neurons are highly specialized cells. Therefore, to better understand how flicker affects neuron-specific functions within VC-ME6 genes (Neuronal enriched module), we used GSVA to compute enrichment scores for 14 neuronal gene sets from our prior work, including synaptic plasticity, signaling, neurotransmission, cytoskeleton, metabolism, and mitochondria. Interestingly, although flicker stimulation suppressed neuronal genes in VC-ME6, flicker led to enrichment of genes annotated for synaptic plasticity, signaling and cytoskeleton. Flicker also suppressed metabolism and mitochondria gene sets within VC-ME6 genes (**Figs. 5E, S5, Table S10)**. Together, this finding suggests that while all frequencies of stimulation suppress genes involved in neuronal transcriptional machinery, genes controlling synaptic plasticity and signaling are enriched by all frequencies of stimulation in VC.

**Figure 5:**
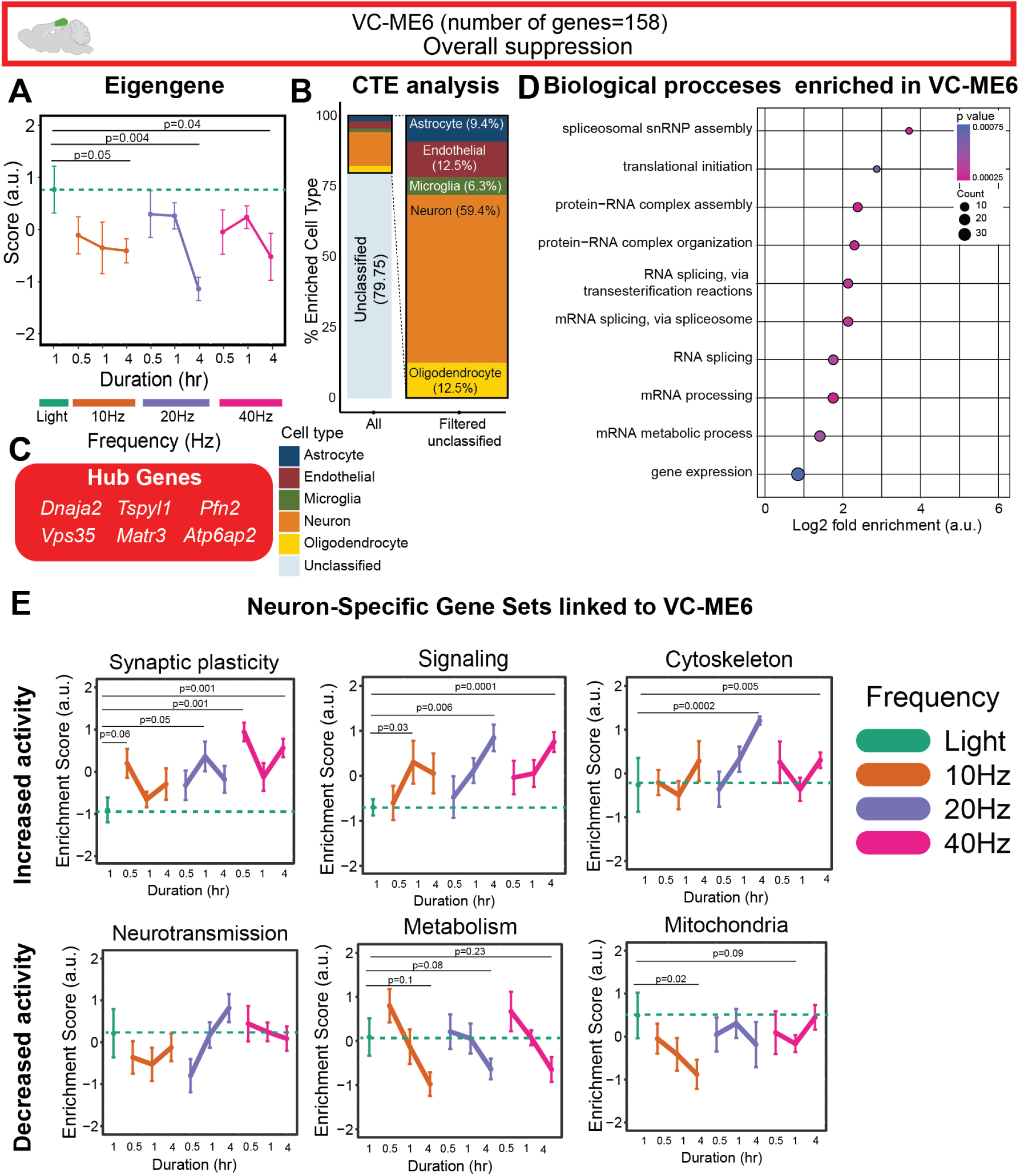
All frequencies of flicker suppress a neuron-enriched gene module and upregulation of neuronal signaling in a duration-dependent manner in visual cortex. (**A**) VC-ME6 (Red) genes were suppressed by all frequencies of flicker in a duration dependent manner (**Table S1**, mean±SEM, linear mixed model). (**B**) Neuronal genes were the most prominent genes enriched in VC-ME6 (**C**) The six most central genes in the module were determined as hub genes of VC-ME6. (**D**) Gene ontology enrichment of biological processes for VC-ME6 (Fisher’s exact test p <0.05) identified transcriptional machinery related genes associated with this module (Dot size represents gene counts and dot colors indicate p-values). (**E**) Gene set variation analysis of VC-ME6 genes using custom cortical neuronal specific gene sets (**Methods**) revealed all frequencies activated of genes associated with synaptic plasticity, signaling and cytoskeleton and suppressed neurotransmission, metabolism, and mitochondrial gene expression in a duration dependent manner.

### Hippocampus exhibits slow suppression of neural differentiation, development and signaling

We next asked if flicker stimulation induced frequency and duration specific effects in the hippocampus, a deep brain region affected early in AD. We conducted a separate WGCNA within the hippocampus (HIP), identifying 15 modules (**Fig. 6, Methods**). Among all hippocampal MEs, (HIP-ME), HIP-ME9, HIP-ME12, HIP-ME1, and HIP-ME10 showed the most pronounced changes in response to flicker duration and frequency (**Fig. S3**). Similar to VC, we excluded the modules with strong correlation from further analysis (**Fig. S6**).

**Figure 6:**
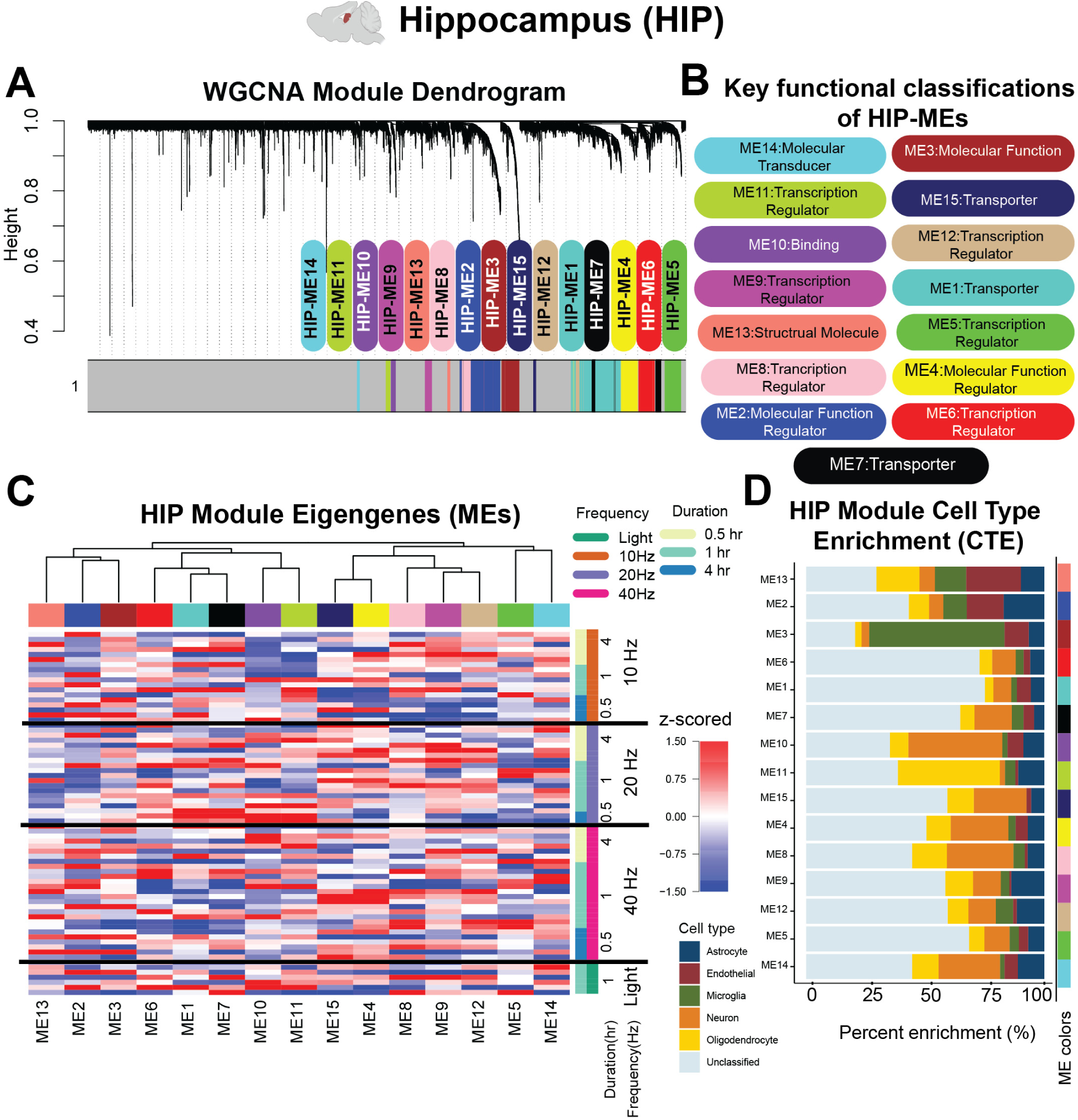
Hippocampus WGCNA identifies modules of co-expressed genes associated with frequency and duration of stimulation. (**A**) WGCNA identifies 15 MEs in HIP. (**B**) Key functional classifications of modules were identified via GO Panther function classification. (**C**) z-scored expression values for 15 MEs in HIP are clustered using Euclidean distance. (**D**) Cell type percent enrichment of all 15 MEs in HIP were calculated by crossing-referencing module genes using GSVA against list of genes previously determined as enriched in neurons, oligodendrocytes, astrocytes, microglia and endothelial. Some MEs are highly enriched by genes expressed by microglia (ME3) and neurons (ME10, ME15, ME4, ME8, and ME14) over other cell-types.

Both HIP-ME9 and HIP-ME12 modules demonstrated slow suppression in response to 10Hz and 20Hz of stimulation. Despite the patterns of HIP-ME9 and HIP-ME12, GO analysis revealed that the genes comprising these modules control different functions. For example, 10Hz or 20Hz frequencies resulted in slow suppression of HIP-ME9, which is enriched for genes associated with behavior, dendritic development, microtubule-based movement, neurotransmitter receptor regulation, and glutamatergic neuron differentiation (**Fig. 7A-D**). HIP-ME12 also showed slow suppression after 10Hz or 20Hz stimulation but was enriched for GO terms involving RTK signaling pathway and response to oxygen species (**Fig. 7H**) where all are dysregulated in AD (39-42). Importantly, neither of these modules demonstrated high specificity to any of five cell types assessed in this paper (**Fig. 7B, G)**. Together, these results show that a single session of prolonged flicker stimulation suppresses multiple hippocampal gene modules, each of which are associated with unique functions with frequency-specific effects.

**Figure 7:**
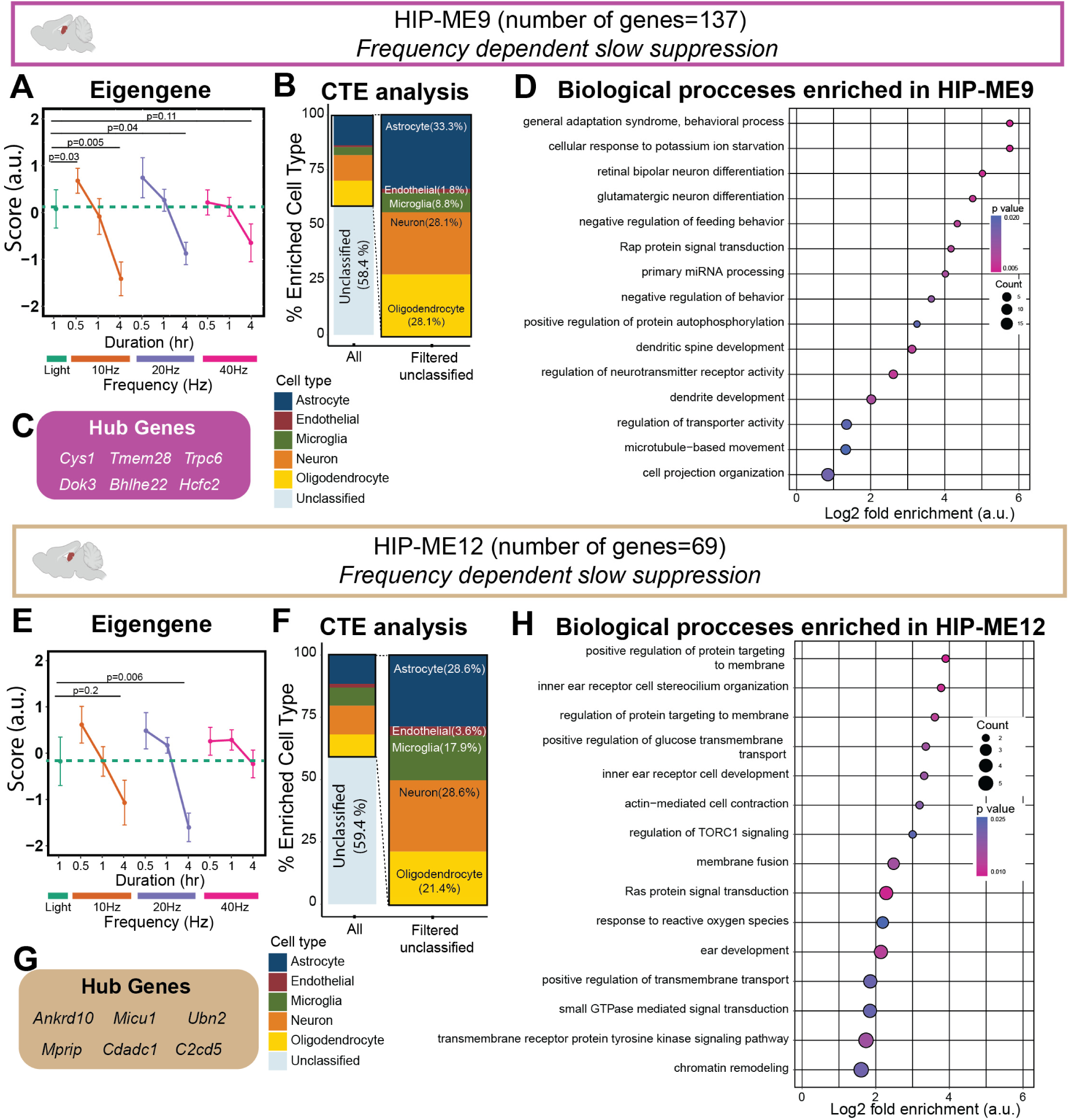
prolonged 10 and 20 Hz flicker modulate neurotransmission and neural differentiation gene expression and activate neuroimmune-signaling genes. (**A**) 10Hz and 20Hz stimulation results in slow suppression of HIP-ME9 (magenta) genes compared to light (**Table S1**, mean±SEM, linear mixed model). (**B**) HIP-ME9 CTE showed enrichment for multiple cell types assessed in this study. (**C**) The six most central genes in the module were determined as hub genes of HIP-ME9. (**D**) Gene ontology enrichment of biological processes for HIP-ME9 genes (Fisher’s exact test p <0.05) identified enrichment of neurotransmission activity, neural differentiation, development, and behavior genes (Dot size represents gene counts and dot colors indicate p-values). (**E**) 10Hz and 20Hz stimulation resulted in slow suppression of HIP-ME12 (tan) genes compared to light (**Table S1**, mean±SEM, linear mixed model). (**F**) CTE showed enrichment for multiple cell types within HIP-ME12. (**G**) The six most central genes in the module were determined as hub genes of HIP-ME12. (**H**) Gene ontology enrichment of biological processes (Fisher’s exact test p <0.05) identified signal transduction and neuroimmune changes associated within HIP-ME12 genes (Dot size represents gene counts and dot colors indicate p-values).

### Hippocampal 10Hz stimulation rapidly suppressed neuronal functions while 20Hz slowly activated mitochondrial metabolism and synapse translation

In addition to the slow suppression effects of HIP-MEs 9 and 12, flicker rapidly suppressed some HIP modules with only 30 min of flicker in a frequency-dependent manner. Specifically, 10Hz flicker rapidly suppressed the neuronally enriched HIP-ME10 (**Fig. 8A, B**). HIP-ME10’s hub genes were also mainly neuronal, including *Fndc9, Slc10a4* and *Sox1,* which are involved in nervous system development, neurotransmitter activity, and neural differentiation, respectively (43-45) (**Fig. 8C**). Although our custom gene sets annotated for cortical neurons (**Fig. 5**) did not reveal significant enrichment in the hippocampus, significant GO terms were associated with neurotransmission activity, neural development, and differentiation and neural signaling in HIP-ME10 (**Fig. 8D**).

**Figure 8:**
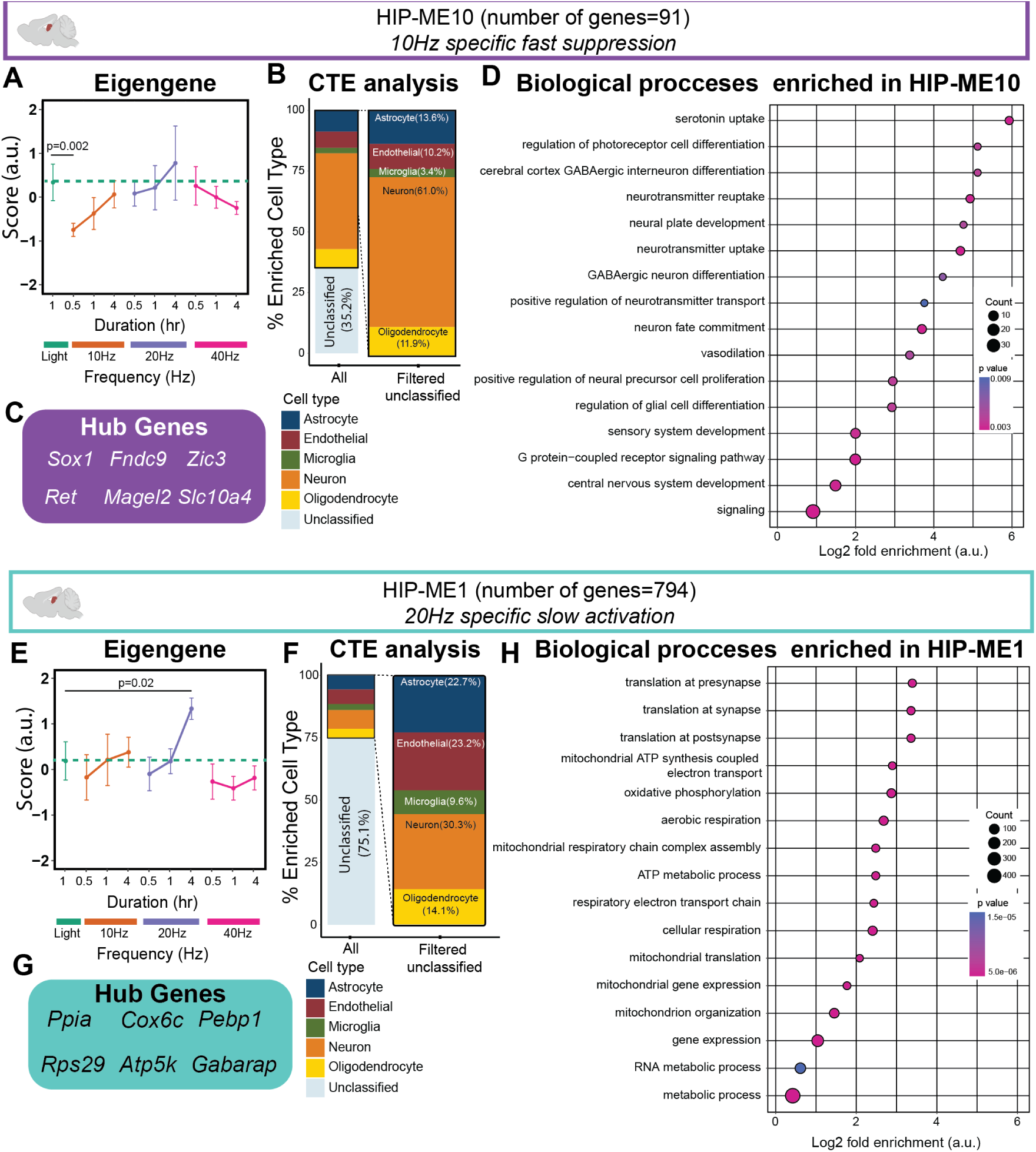
10Hz flicker rapidly suppresses module related to neurotransmission and differentiation while 20Hz slowly stimulates module of mitochondrial and energy source genes. (**A**) 10Hz stimulation rapidly suppressed HIP-ME10 (purple) genes compared to light (**Table S1**, mean±SEM, linear mixed model). (**B**) Neuronal genes were the most prominent genes enriched in HIP-ME10 (∼60%). (**C**) The six most central genes in the module were determined as hub genes of HIP-ME10. (**D**) Gene ontology enrichment of biological processes for HIP-ME10 genes (Fisher’s exact test p <0.05) identified neurotransmitter regulation and neural differentiation as the main functions driven by genes in HIP-ME10 (Dot size represents gene counts and dot colors indicate p-values). (**E**) 20Hz stimulation slowly activated HIP-ME1 (turquoise) genes relative to light (**Table S1**, mean±SEM, linear mixed model). (**F**) CTE analysis showed enrichment of multiple cell types assessed in this study. (**G**) The six most central genes in the module were determined as hub genes of HIP-ME1. (**H**) Gene ontology enrichment of biological processes for ME1 genes (Fisher’s exact test p <0.05) identified synaptic translation, metabolism, and mitochondrial regulatory processes associated with the genes in HIP-ME1 (Dot size represents gene counts and dot colors indicate p-values).

Flicker also induced frequency-specific activation in HIP. Specifically, 20Hz flicker caused slow activation of HIP-ME1, which was enriched for GO terms associated with mitochondrial function, gene expression, and translation at synapses (**Fig. 8E-8H**). Thus, flicker can be tuned based on frequency and duration to activate or suppress a distinct set of genes in the hippocampus.

Although the HIP modules changing in response to flicker stimulation are distinct from VC, gene annotations in modules from both regions share many functions, including neurotransmission, dendrites remodeling, and neuro-immunity (**Fig. 9**). Nevertheless, different frequencies of stimulation had specific effects in each region. We found that 40Hz drives transcriptional changes linked to synaptic activity, neuronal function such as neurotransmission activity and neuron remodeling (VC-ME9, VC-ME5), and gene expression (VC-ME6) in VC, but did not identify any significant changes driven by 40Hz stimulation in HIP. Also in VC, prolonged 20Hz stimulation suppressed neuronal transcriptional machinery (VC-ME6), while in HIP, the same frequency and duration suppressed genes associated with neuroimmune signaling (HIP-ME12) and activated metabolism and synaptic translation (HIP-ME1). Furthermore, 10Hz stimulation led to rapid activation of neuroimmune signaling in visual cortex (VC-ME8), but rapid suppression of neurotransmitter activity and neuronal differentiation in hippocampus (HIP-ME10). Thus, our data collectively indicate that, while similar effects can be achieved in VC and HIP, the frequencies and durations of stimulation need to be tuned based on both the cellular function and the region of the brain being targeted.

**Figure 9:**
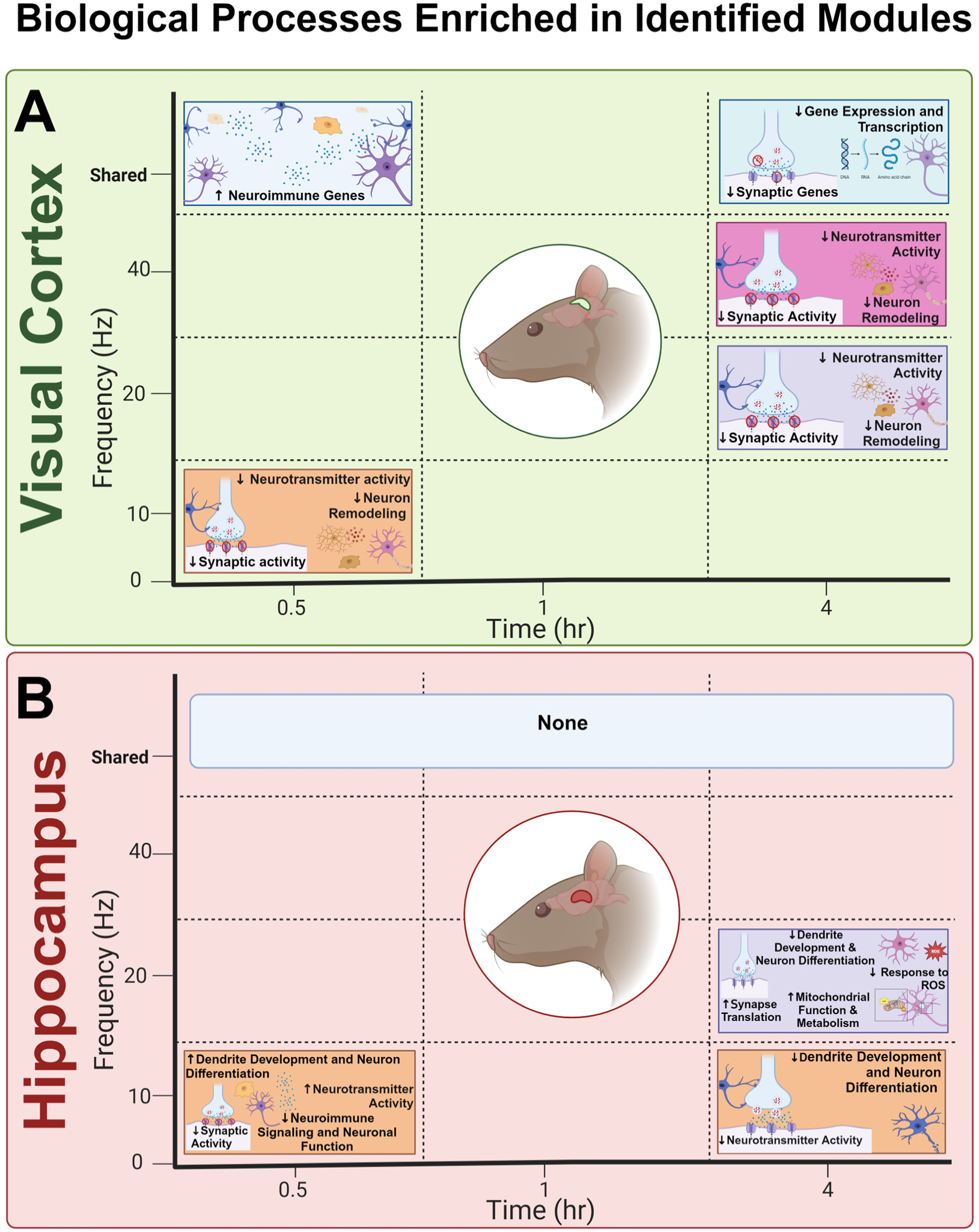
Map of stimulation parameters into transcriptional changes in the visual cortex and hippocampus. **(A)** Within the VC, flicker stimulation drives neuroimmune, neuronal, and synaptic changes with patterns that are either shared across all frequencies or specific to certain ones. The shared changes are associated with upregulation of neuroimmune genes at 0.5hr and reduced synaptic function and gene expression at 4hr. The distinctive changes include reduced neurotransmission neural remodeling after either 0.5hr of 10Hz or 4hr of 20Hz stimulation. (**B**) Within the HIP, flicker did not elicit shared changes across frequencies at any time point. At 0.5hr of 10Hz stimulation, dendrite development, neurotransmitter activity, and neuronal differentiation processes were elevated, but similar processes were decreased after 4hr of 10Hz. Additionally, 4hr of 20Hz stimulation led to increased synapse translation, reduced dendrite development, and increased mitochondrial function and metabolism.

## Discussion

This study presents the most detailed analysis to date of the multipotent effects of multiple frequencies and durations of flicker stimulation in two brain regions of the 5xFAD amyloid mouse model of Alzheimer’s disease. We hypothesized that distinct frequencies and durations of stimulation would differentially modulate gene expression relevant to the functions of neurons, astrocytes, and microglia. Because 5xFAD mice develop cognitive deficits starting at 4mo (46, 47), we used 3-7mo animals to evaluate the effects of flicker in the early symptomatic age range, which is relevant to clinical therapy. By simultaneously analyzing the effects of frequency and duration of flicker, this is the first study to pinpoint stimulation parameters (i.e., frequency and duration) that optimally modulate transcriptional signatures associated with either neuronal or glial functions. Our data highlights that the optimal stimulation protocol is determined based on the brain process and region to be targeted (**Fig 9**).

Two key stimulation variables we examined in this study are frequency and duration. We used WGCNA to provide an unbiased summary of gene expression trends in our data, then assessed each identified gene module for enhanced expression of specific biological processes (gene ontology) and cell type specific enrichment (CTE). Indeed, we identified a shared effect of frequency in VC, where all frequencies (10, 20 and 40Hz) led to fast activation neuroimmune genes (VC-ME8), prolonged regulation synaptic activity and plasticity (VC-ME5) and prolonged suppression of neuronal transcriptional machinery (VC-ME6). Within HIP, we did not observe any shared effects among different frequencies and many of the flicker-induced effects were frequency specific. For example, 10Hz stimulation acutely suppressed modules of genes associated with neuron fate commitment and neurotransmitter uptake (HIP-ME10), while 20Hz stimulation led to prolonged activation of synaptic translation, mitochondrial function, and metabolism (HIP-ME1). We found some of the modules effected by flicker stimulation across both brain regions were cell type specific. Specifically, genes in VC-ME9 linked to synaptic pruning and activity were enriched in astrocyte-specific genes (∼39%) while genes in VC-ME6 linked to transcriptional machinery and genes in HIP-ME10 linked to neurotransmitter activity were profoundly enriched in neurons.

An extensive body of prior work has investigated the effects of flicker in response to 1hr of stimulation. Prior studies have shown that 40Hz flicker has multiple effects, including reducing amyloid beta levels, altering microglia and astrocyte morphology, increasing glymphatic clearance, and more (6-9, 17). These prior studies primarily focused on the effects of 1hr of 40Hz flicker or repeated exposures to 1hr of 40Hz flicker. Additionally, our own prior work found that that short durations of flicker, as little as 5min, resulted in activation of MAPK pathway signaling, which is known to affect downstream transcription in neuronal, synaptic, and immune pathways in a duration-dependent manner (48, 49). Consistent with the known duration-dependence of MAPK and NFκB, we found that different durations of the same frequency elicited different downstream transcriptional changes (**Fig. 9**). Within HIP, short duration (30min) of 10Hz stimulation activated modules of genes linked to dendrite development and neurotransmitter activity, while longer duration of 10Hz led to suppression of a distinct set of genes which surprisingly linked to similar function. In other cases, the different durations led to differential expression of genes with distinction functional roles. For example, within VC, while prolonged (4hr) 20Hz stimulation led to reduction in expression of module of genes linked to synaptic activity, 1hr of 20Hz led to activation of module of genes linked to neuroimmune activity. In addition, 1hr of 10Hz stimulation led to suppression of neurotransmission activity and synaptic function within astrocyte (VC-ME9) while 4hr of 10Hz stimulation led to suppression of neuronal transcriptional machinery (VC-ME6). Together, these findings emphasize the importance of duration of stimulation in defining upstream signaling pathways and downstream transcriptional changes.

A key finding from our study is that audiovisual flicker affects HIP in addition to the VC, which is clinically important to target in Alzheimer’s disease. While prior studies have reported that 20 and 40Hz visual stimulation for 1hr could induce molecular changes within the visual cortex, it was previously unknown whether or not just one of 1hr of audiovisual stimulation could module the HIP transcription profile. While our data show that audiovisual flicker effects both regions, we found that the specific transcriptional changes are highly dependent on the region being targeted. Nevertheless, despite our observed differences in the two brain regions, we showed that each frequency or duration of audiovisual flicker led to specific transcriptional changes in each region.

This study has several important limitations that necessitate future work. First, we limited the current work to males, due to the known sexual dimorphism in 5xFAD mice and the need to be sufficiently powered across multiple frequencies and durations of stimulation. Second, we relied on bulk RNAseq to enable us to quantify transcriptomes across a total of 177 samples in two brain regions and two genotypes, but many of the pathways we identified are central regulators of cellular function in neurons, microglia, and other cell types. Our data support the effects of flicker in multiple major cell types and identify transcriptional changes associated with specific cell-types. In future studies, single cell, or single nucleus RNAseq will be essential to uncover how these pathways are becoming activated in specific cell types. Third, we used mRNA quantification to define flicker-induced changes in 5xFAD mouse brain, but there is limited concordance between mRNA and protein-level changes. Therefore, proteomics and functional assays will be essential to assess the effect of flicker stimulation most accurately. Fourth, mice selected for this study had an age range from 3-7mo old, due to collection constraints during the pandemic. To ensure the age of the mice was not a driver of the transcriptional signatures we found here, we correlated age all identified modules (**Fig. S6**) and excluded any that were significantly correlated. Nevertheless, because we found differences in the flicker induced transcriptional profiles in 5xFAD and WT mice (**Fig. 1**), its effects will need to be evaluated in future studies at detailed timepoints during onset and progression of Alzheimer’s-like pathology.

## Conclusion

In summary, we found that transcriptional signatures, e.g., VC-ME8, VC-ME5, and VC-ME6 are shared across all frequencies of stimulation, whereas others, e.g., VC-ME9, VC-ME6, HIP-ME10, and HIP-ME1 are specific to the frequency as well as the duration. The results highlight the importance of frequency, duration, and region in defining the effects of audiovisual stimulation in the context of AD-pathology. We also demonstrated that gene modules targeted by flicker are linked to multiple functions that are affected in AD. Moreover, we demonstrated that different frequencies and durations modulate multiple biological functions and cell types, which opens new avenues for novel non-invasive and multi-potent treatment of AD. Collectively, our data indicate that the frequency and duration of flicker stimulation controls immune, neuronal, and metabolic genes in multiple regions of the brain affected by Alzheimer’s disease. Flicker stimulation may thus represent a potential therapeutic strategy that can be tuned based on the brain region and the specific cellular process to be modulated.

## Supporting information

Supplemental Table 2_DEGs_5xFAD_1hr_20 vs 40 Hz

Supplemental Table 3_Go terms associated with DEGs 5xFAD 40Hz vs 20Hz_VC and HIP

Supplemental Table 4_DEGs WT 40Hz vs 20Hz_VC and HIP

Supplemental Table 5_ GO terms associated with Modules in VC

Supplemental Table 6_ GO terms associated with Modules in HIP

Supplemental Table 7_DEGs 5xFAD Light vs 10Hz 20Hz 40Hz _1hr duration _VC and HIP

Supplemental Table 8-summary of ME stats using linear mixed model

Supplemental Table 9-summary of VC_ME9 Custom GS_ Stats using linear mixed model

Supplemental Table 10-summary of VC_ME6 Custom GS_ Stats using linear mixed model

Supplemental Table 11-RRID

## Availability of Data and Materials

The gene expression FASTQ files and count matrix that support the findings of this study have been deposited in the Gene Expression Omnibus (GEO).

## Author Information

### Authors and Affiliations

*George W. Woodruff School of Mechanical Engineering, Georgia Institute of Technology*

Sara Bitarafan, Felix G. Rivera Moctezuma, Levi B. Wood

*Parker H. Petit Institute for Bioengineering, Georgia Institute of Technology*

Sara Bitarafan, Alyssa F. Pybus, Felix G. Rivera Moctezuma, Levi B. Wood

*Wallace H. Coulter Department of Biomedical Engineering, Emory University and Georgia Institute of Technology*

Alyssa F. Pybus, Tina C. Franklin, Annabelle C. Singer

*School of Biological Sciences, Georgia Institute of Technology*

Mohammad Adibi

### Contributions

LBW and ACS designed and oversaw execution of the study. SB, AFP, MA, and LBW conducted experiments. SB and FGR prepared figures. LBW and SB designed the computational strategy and conducted data analysis. TCF advised on methods and interpretation of results. SB, LBW, ACS drafted the manuscript. All authors critically reviewed the data analysis and manuscript.

### Corresponding Authors

Correspondence to Annabelle C. Singer and Levi B. Wood.

## Funding

This work was supported by funds from The Rotary Coins for Alzheimer’s Research Trust Fund (LBW/ACS). L.B.W. acknowledges funding from the National Science Foundation under award number CAREER 1944053, the National Institutes of Health under grant number NIH R01AG075820, and the George W. Woodruff School of Mechanical Engineering. A.C.S. acknowledges the Packard Foundation, NIH NINDS R01 NS109226, NIH NINDS RF1NS109226, NIH NIA RF1AG078736-01, Bright Focus Foundation Grant A2022048S, the McCamish Foundation, and the Lane Family.

## Acknowledgment

We wish to thank the core facilities at the Parker H. Petit Institute for Bioengineering and Bioscience at the Georgia Institute of Technology for use of their shared equipment, services, and expertise.

## Ethics Declarations

### Ethics approval and consent to participate

All protocols were approved by Georgia Institute of Technology Institutional Animal Care and use Committee (IACUC) in accordance with National Institutes of Health guidelines.

### Consent for publication

Not applicable.

### Competing interests

The authors have no conflicts of interest to disclose.

## Additional Figures and Tables

**Supplementary Table S1: Sample size of each experimental group.**

**Supplementary Table S2: Differentially expressed genes between 1hr of 20Hz and 40Hz stimulation in 5xFAD mice.**

**Supplementary Table S3: Go terms associated with DEGs between 40Hz vs 20Hz of 5xFAD mice across both regions.**

**Supplementary Table S4: Differentially expressed genes between 1hr of 20Hz and 40Hz stimulation in WT mice across both regions.**

**Supplementary Table S5: GO terms associated with VC MEs**

**Supplementary Table S6: GO terms associated with HIP MEs**

**Supplementary Table S7: Differentially expressed genes between 1hr of 10, 20, or 40Hz of flicker in 5xFAD mice across both regions.**

**Supplementary Table S8: Summary of ME stats using linear mixed model.**

**Supplementary Table S9: summary of VC_ME9 Custom GS_ Stats using linear mixed model.**

**Supplementary Table S10: summary of VC_ME6 Custom GS_ Stats using linear mixed model.**

**Supplementary Table S11: RRIDs of resources.**

**Supplementary Figure S1: Differentially expressed genes between 1hr of 20Hz and 40Hz stimulation in WT mice.**

**Supplementary Figure S2: WGCNA identifies 12 Module Eigengenes (MEs) in VC.**

**Supplementary Figure S3: WGCNA identifies 15 Module Eigengenes (MEs) in HIP.**

**Supplementary Figure S4: Custom astrocyte specific gene sets associated with VC-ME9.**

**Supplementary Figure S5: Custom neuron specific gene sets associated with VC-ME6.**

**Supplementary Figure S6: Correlation heatmap between MEs and age.**

## Notes

### Competing Interest Statement

The authors have declared no competing interest.

