## Supplementary Figures and Supplemental Table 1 for "Frequency and duration of sensory flicker controls astrocyte and neuron specific transcriptional profiles in 5xFAD mice"

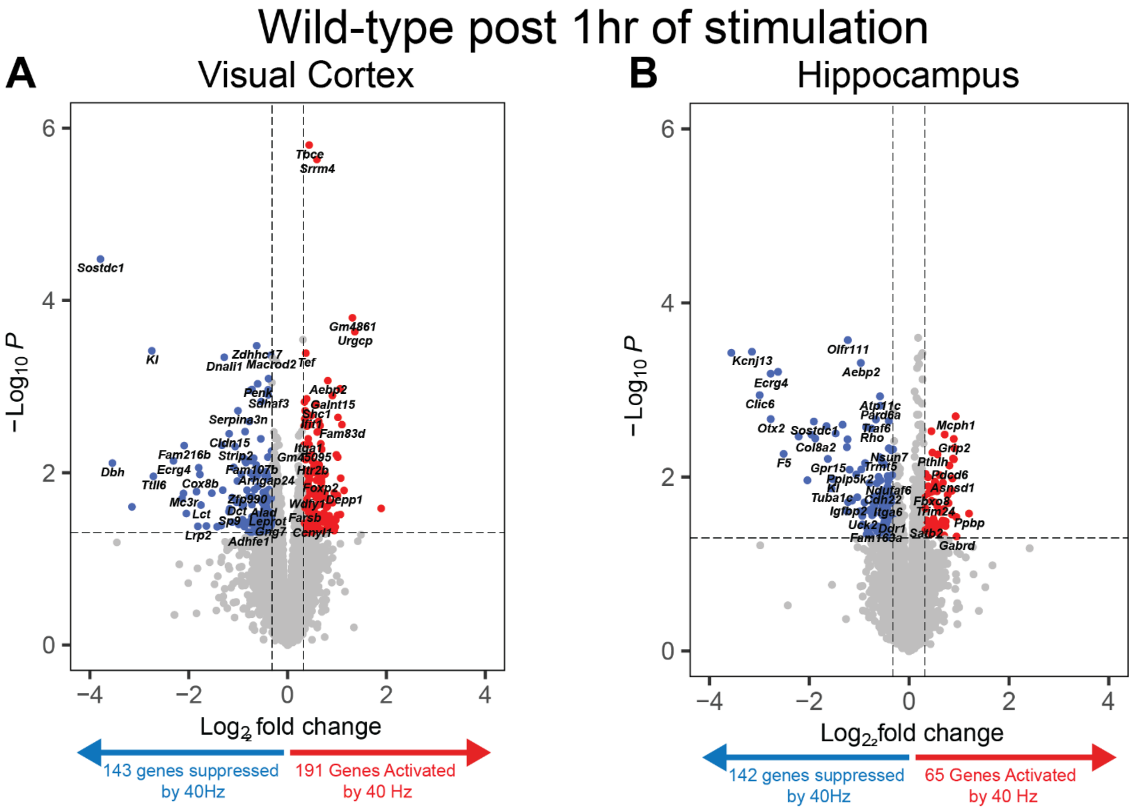


**Figure S1: Differentially expressed genes between 1hr of 20Hz and 40Hz stimulation in WT mice.** (**A**) 40Hz vs 20Hz stimulation in the visual cortex resulted in differentially expressed genes after 1hr (|$\log_2$ fold change| > 0.25, unadjusted p<0.05). (**B**) 40Hz vs 20Hz stimulation in the hippocampus resulted in differentially expressed genes after 1hr (|log2 fold change| > 0.25, p<0.05).

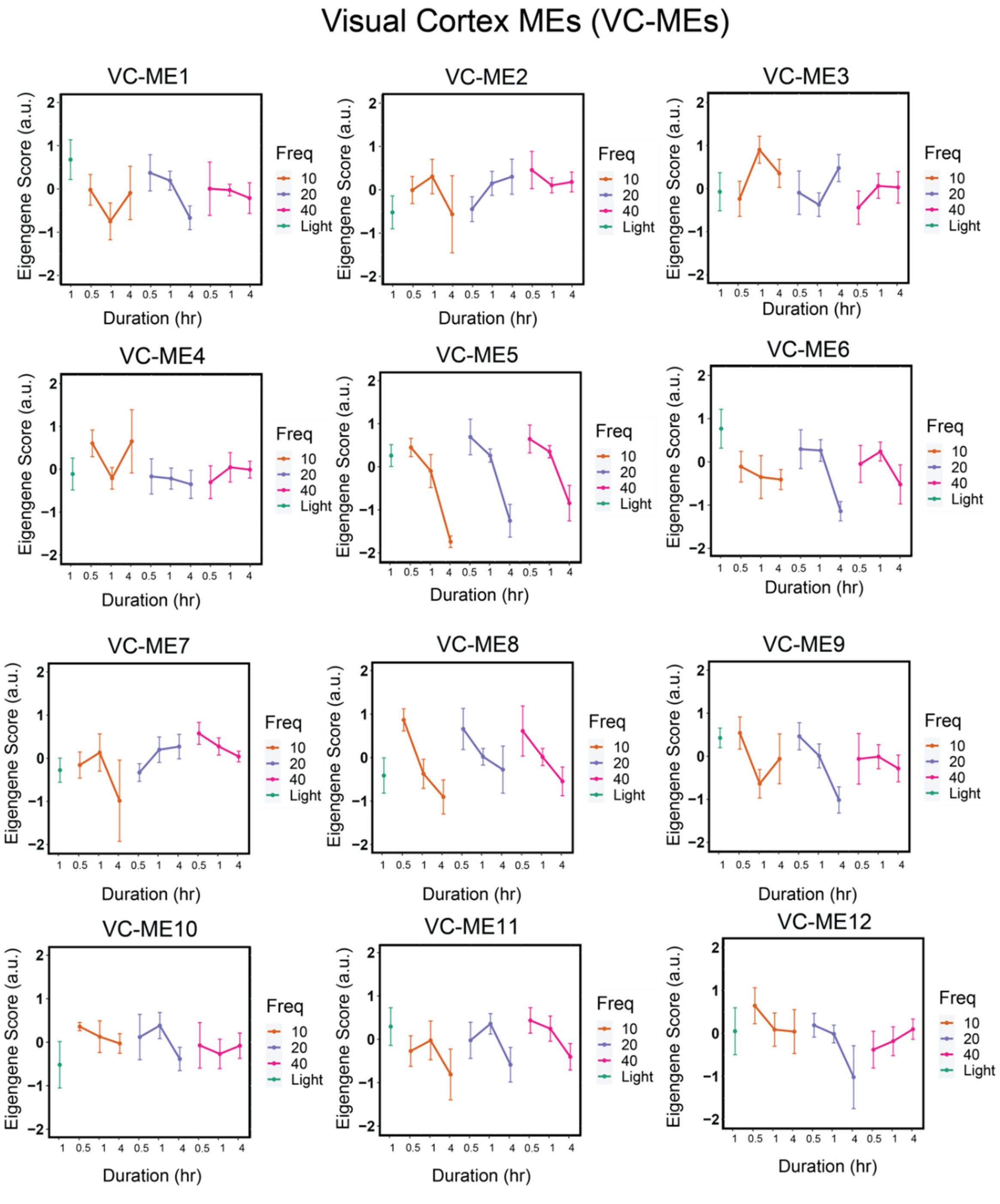


**Figure S2: WGCNA identifies 12 Module Eigengenes (MEs) in VC.** VC-MEs demonstrated distinct frequency-duration dependent response to flicker stimulation (See **Table S1** for sample size per group, mean±SEM).

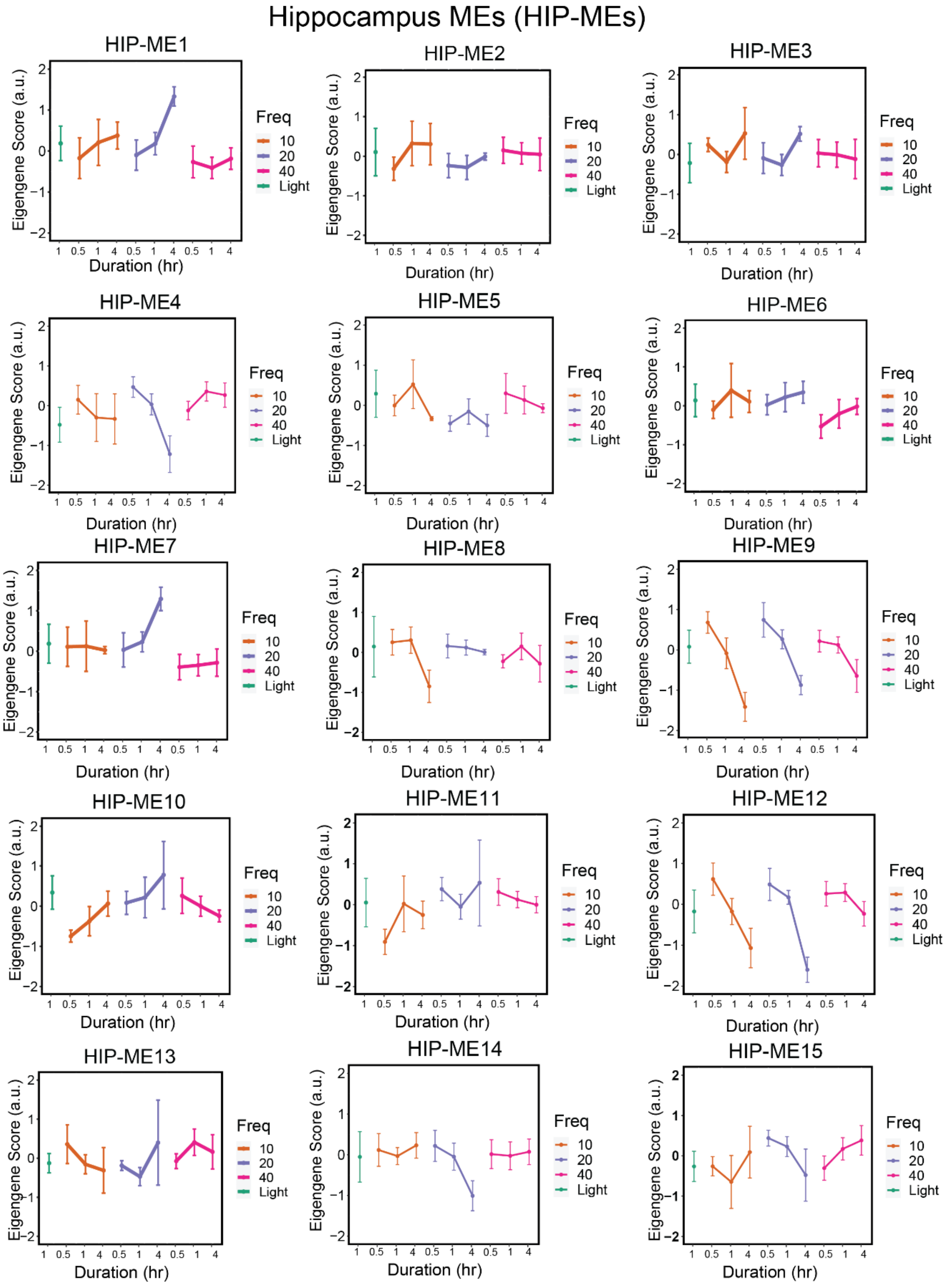


**Figure S3: WGCNA identified 15 Module Eigengenes (MEs) in HIP.** HIP-MEs demonstrated distinct frequency-duration dependent response to flicker stimulation (See **Table S1** for sample size per group, mean±SEM).

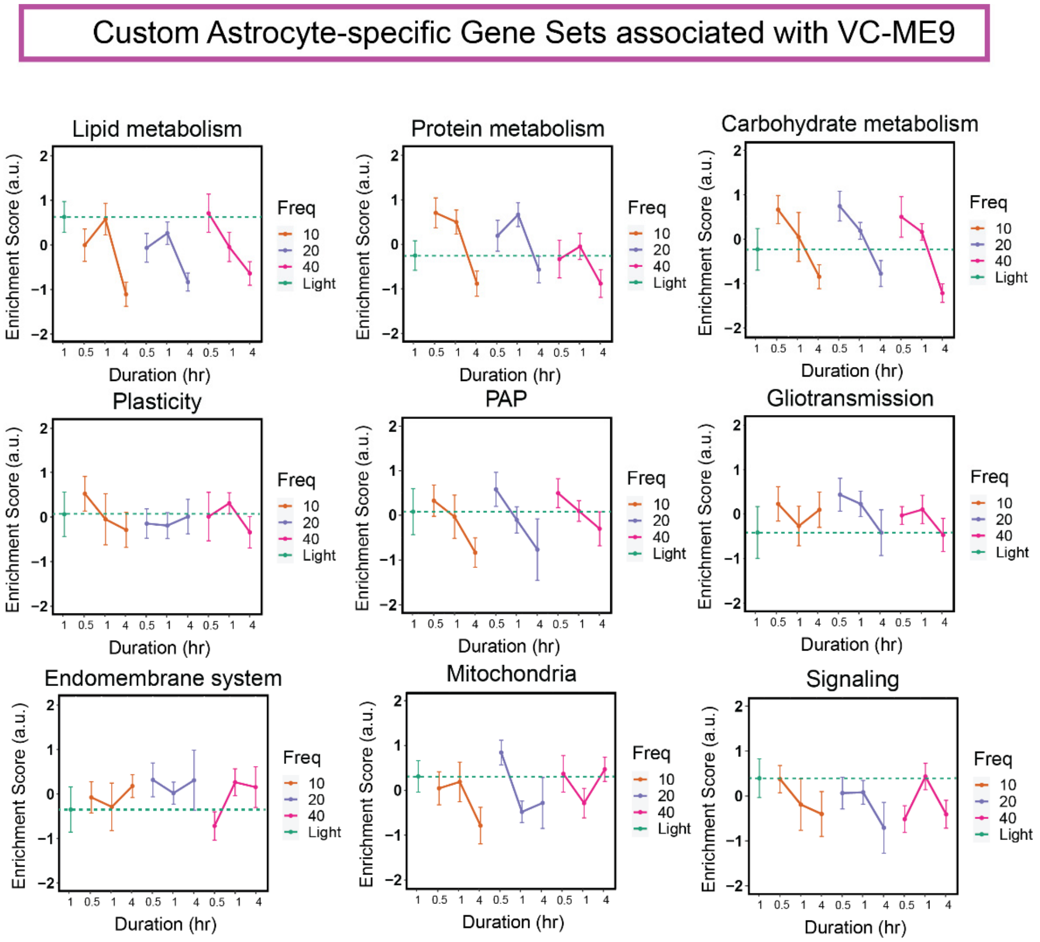


**Figure S4: Enriched Custom astrocyte specific gene sets associated with VC-ME9 identified by GSVA.** Custom astrocyte gene set variation analysis of genes in VC-ME9 module demonstrated distinct pattern of astrocytic function related gene sets in response to different frequencies and durations of flicker stimulation (See **Table S1** for sample size per group, mean±SEM).

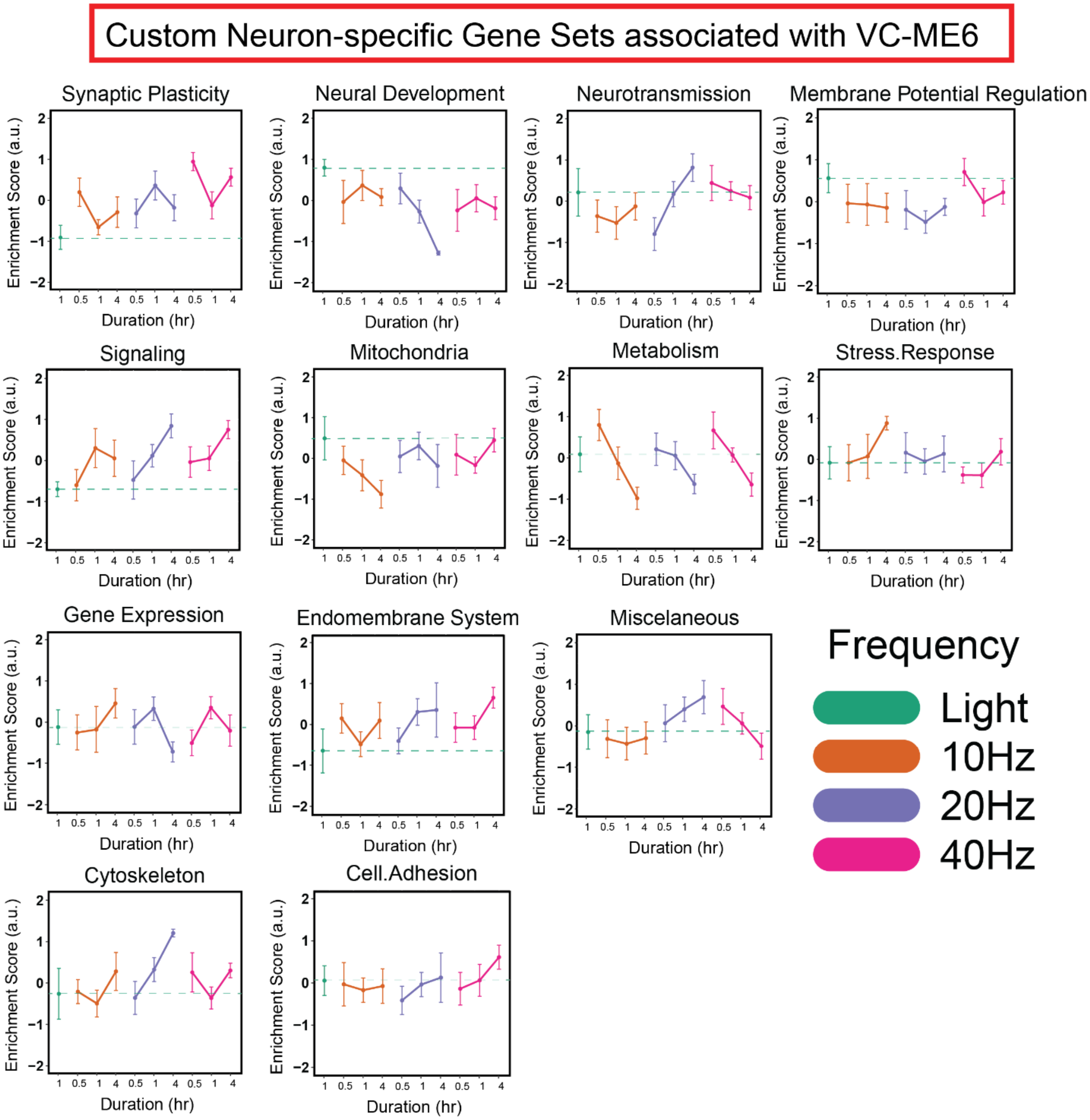


**Figure S5: Enriched Custom neuron specific gene sets associated with VC-ME6 identified by GSVA.** Custom neuronal gene set variation analysis of genes in VC-ME6 module demonstrated distinct pattern of neuronal function related gene sets in response to different frequencies and durations of flicker stimulation (See **Table S1** for sample size per group, mean±SEM).

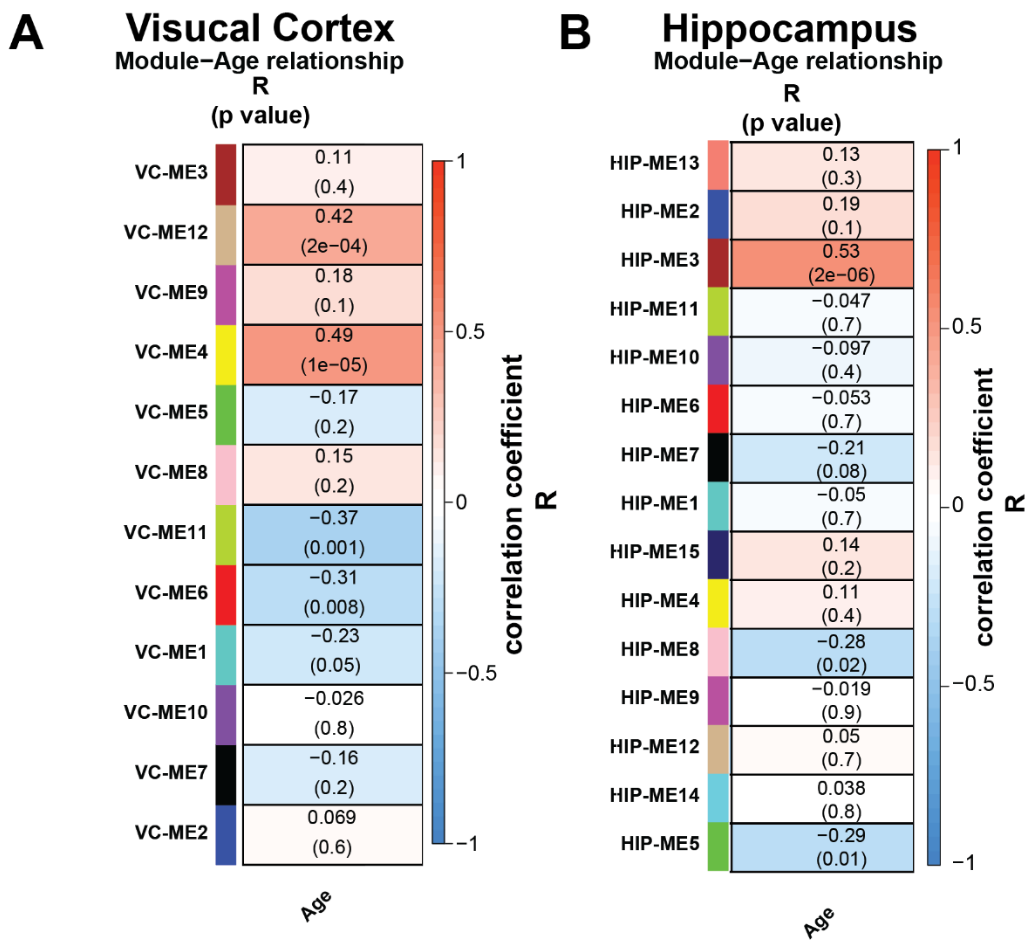


**Figure S6: Correlation heatmap between MEs and age.** Heatmap of Pearson's correlation coefficient of mouse age versus module eigengene (MEs) (rows). The color bar represents correlation coefficient where negative -1 represents strong negative correlation (dark blue) and +1 represents strong positive correlation (dark red). p values associate with slope are indicated in prentices.

### Supplementary Tables:

**Table S1: Sample size of each experimental group.** Sample size (N, left column) for each genotype, brain region, flicker frequency, and stimulation duration.

| Genotype | Region | Frequency | Duration | N |
|---|---|---|---|---|
| 5xFAD | VC | Light | 1hr | 6 |
| | | 10Hz | 0.5hr | 7 |
| | | | 1hr | 6 |
| | | | 4hr | 6 |
| | | 20Hz | 0.5hr | 7 |
| | | | 1hr | 12 |
| | | | 4hr | 3 |
| | | 40Hz | 0.5hr | 6 |
| | | | 1hr | 13 |
| | | | 4hr | 7 |
| | HIP | Light | 1hr | 6 |
| | | 10Hz | 0.5hr | 7 |
| | | | 1hr | 6 |
| | | | 4hr | 6 |
| | | 20Hz | 0.5hr | 7 |
| | | | 1hr | 12 |
| | | | 4hr | 3 |
| | | 40Hz | 0.5hr | 7 |
| | | | 1hr | 13 |
| | | | 4hr | 7 |
| WT | VC | 20Hz | 1hr | 8 |
| | | 40Hz | | 7 |
| | HIP | 20Hz | 1hr | 8 |
| | | 40Hz | | 7 |
